# Synovial macrophage activation mediates pain experiences in experimental knee osteoarthritis

**DOI:** 10.1101/2023.04.25.538261

**Authors:** Garth Blackler, Yue Lai-Zhao, Joseph Klapak, Holly T. Philpott, Kyle K. Pitchers, Andrew R. Maher, Benoit Fiset, Logan A. Walsh, Elizabeth R. Gillies, C. Thomas Appleton

## Abstract

It has been suggested that synovial macrophages mediate nociceptive signals in knee osteoarthritis (OA) but the underlying mechanisms are unknown. Our objectives were to investigate the role of synovial macrophages and their activation via signal transducer and activator of transcription (STAT) signaling in mediating OA pain experiences.

We induced experimental OA in rats via knee destabilization surgery and then performed RNA sequencing analysis in sorted synovial macrophages to identify signaling pathways associated with macrophage activation. Next, we repeated intra-articular injections of liposomal clodronate to deplete macrophages, or liposomal inhibitors of STAT1 or STAT6 to block macrophage activation, and tested the effects on local and distal mechanical pain sensitivity. We also assessed synovitis, cartilage damage, and synovial macrophage infiltration with histopathology and immunofluorescence, and crosstalk between liposomal drug-treated synovium and articular chondrocytes in co-culture.

Most enriched signaling pathways in activated OA macrophages involved STAT signalling. Macrophage depletion and STAT6 inhibition led to marked, sustained improvements in mechanical pain sensitivity and synovial inflammation compared to controls, but macrophage depletion caused increased synovial fibrosis and vascularization. In contrast, STAT1 and STAT6 inhibition in macrophages did not worsen synovial or cartilage pathology. In crosstalk assays, macrophage STAT1-inhibited synovium caused the greatest increases in the expression of anabolic and catabolic chondrocyte genes and sulphated glycosaminoglycan secretion in chondrocytes.

Our results suggest that synovial macrophages play a key role in mediating pain experiences in experimental knee OA, and that selectively blocking STAT6 in synovial macrophages may reduce OA-related pain without accelerating joint tissue damage. (248/250)

**One Sentence Summary:** Selective drug targeting to synovial macrophages improves pain experiences in surgical joint destabilization-induced experimental rodent knee OA. (145/150)

## Introduction

Available treatments for chronic joint pain in knee osteoarthritis (OA) are limited by off-target effects. Recently, a biologic anti-nerve growth factor (NGF) (tanezumab) successfully modified pain but was rejected by regulators due to a small but important increase in rapidly progressive OA (*1*). There is an urgent need to identify more selective targets to modify OA-related pain without accelerating joint tissue damage. Synovial joint inflammation (synovitis) is dominated by macrophage infiltration and activation, and strongly associated with worse pain and joint damage progression in patients with knee OA (*2–4*). We and others suspect that synovial macrophages play important roles in mediating pain outcomes, but the mechanisms controlling synovial macrophage activation in OA are not well understood.

Macrophages represent the dominant immune cell in healthy synovium and maintain tissue homeostasis through specialized functions including phagocytosis and efferocytosis to remove by-products of tissue turnover and dead/dying cells, respectively (*5, 6*). During OA development and progression, peripheral macrophages are recruited to synovial tissue and chronically activated (*7*). Macrophages may play a pathogenic role in chronic OA-related pain directly through interactions with nociceptive fibers and/or indirectly with other cell types (*8*). However, the role of macrophages in mediating OA-related pain is not clearly understood.

Due to their plasticity, macrophages exhibit a wide range of activation states through exposure to disease-specific cues in the local joint microenvironment (*9*). Macrophage activation and polarization has been modeled *in vitro* as pro-inflammatory (“M1-like”, producing catabolic factors such as TNF-α), or anti-inflammatory (“M2-like”, producing anabolic factors such as TGF-β, and function in tissue repair) (*10*). However, the M1/M2 paradigm is not disease-specific and may not accurately reflect synovial macrophage activation mechanisms in OA. Identifying disease-specific macrophage activation profiles is therefore required to link activation mechanisms to the role of macrophages in mediating OA-related pain experiences (*5*).

Intra-cellular signaling mechanisms mediate environmental cues to activate macrophages (*9*). Such mechanisms are an attractive target for treating OA-related pain given the success of this strategy in treating other forms of arthritis, including rheumatoid and psoriatic arthritis with Janus kinase (JAK) inhibitors (*11*). We therefore require a better understanding of which intra-cellular pathways are involved in synovial macrophage activation during the development of OA-related pain. Transcription factors such as the STAT (signal transducer and activator of transcription) family play major roles in controlling macrophage activation and polarization in other contexts through direct transcriptional programs and via crosstalk with other signaling pathways (*9, 12*). STAT signaling mechanisms may therefore be master regulators of macrophage activation and could be exploited therapeutically to reduce OA-related pain. Further, we can directly study the effects of STAT inhibitors on phagocytes *in vivo*, while avoiding off-target effects in other cell types, by encapsulating small molecule STAT inhibitors in large multilamellar liposomes for intra-articular injection in small animal models of OA (*13*). Surgically-induced destabilization of the knee joint via anterior cruciate ligament transection (ACLT) and/or destabilisation of the medial meniscus (DMM) in rats and mice are the most commonly used small animal models of human knee OA due to the ability to control the timing of OA onset and well-established pain outcome measures (*14*).

Our objectives were to identify intra-cellular signaling pathways involved in synovial macrophage activation during early OA development and progression, investigate the roles of synovial macrophage activation via STAT1 and STAT6 signaling in mediating OA-related pain, and explore the effects of STAT inhibition on joint histopathology and crosstalk between synovium and articular chondrocytes.

## Results

### Differential gene expression and gene set enrichment analysis

Forty-two genes were differentially expressed in synovial macrophages at 4-weeks after OA induction (18 increased and 24 decreased) (Sup. 1). These included genes involved in cellular metabolism (*Fbp1, Acod1, C1qtnf3, Slc1A3)*, macrophage activation (*Rnase2, Ocstamp*), inflammation (*Sctr*), cell motility (*Cdh26*), and Wnt signalling (*Rspo2)* (Fig. 1A). One hundred thirty-three genes were differentially expressed in synovial macrophages at 12-weeks (78 increased and 55 decreased) (Sup. 2). These included genes involved in extracellular remodelling (*Mmp16, Fbn2, Adamts16, Mmp13, Vit, Cemip*), cell motility (*Myh7, Tnni1*), and cell fate (*Ptprv*) (Fig. 1B).

**Figure 1.**
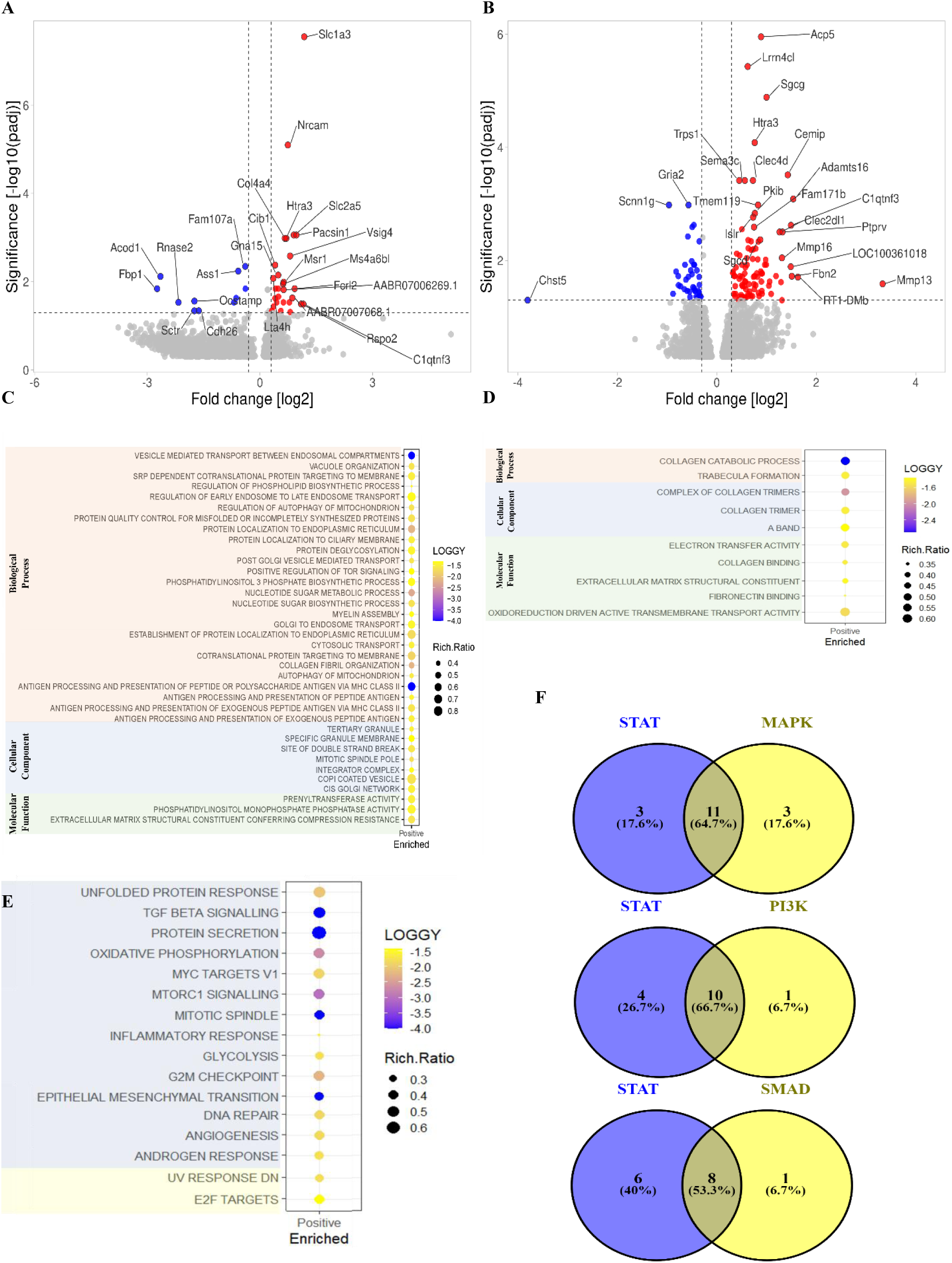
Differential gene expression and gene set enrichment analysis of OA macrophages. Volcano plots showing the top differentially expressed genes of macrophages at 4-weeks **(A)** and 12-weeks **(B)** post surgery, as compared to sham controls. The Y-axis represents the -log10 of the adjusted p-value with a cut-off set at 1.3 (padj < 0.05) and the x-axis represents the log2 fold change with cut-off at −0.5 and 0.5. Bubble plots showing gene set enrichment of Gene Ontology terms at 4-weeks **(C)** and 12-weeks **(D)** and enrichment of Hallmark term at 4-weeks **(E)** after OA induction versus sham controls. LOGGY represents the - log10 of the false discovery rate (FDR <0.05) and Rich.Ratio the number of genes called to a set divided by the total number of genes in the set. Venn diagrams comparing the number significantly enriched hallmark pathways in early-stage OA development (shown in panel E) that are associated with STAT signaling and their overlap with pathways involving MAPK, PI3K, and SMAD signaling mechanisms **(F)**.

Gene ontology analysis found 46 gene sets that were positively enriched at 4 weeks (OA vs sham) (Sup. 3), including cellular metabolism, cellular stress, protein secretion, and macrophage function (Fig. 1C). Interestingly, gene ontology in macrophages from 12-weeks post-surgery (OA vs sham) demonstrated that most enriched gene sets were predominately involved in extracellular matrix remodeling (Fig. 1D, Sup. 4). Hallmark gene set enrichment revealed 16 significantly enriched pathways associated with synovial macrophage activation at 4 weeks post-OA induction (Fig. 1E, Sup. 5). Intracellular signaling pathways involved in the significantly enriched pathways included STAT (14/16 pathways), mitogen activated protein kinase (MAPK; 14/16), phosphoinositide 3-kinase (PI3K; 11/16), and SMAD signaling (9/16; Fig. 1E, Sup. 5). Interestingly, STAT signaling-related pathways overlapped substantially with MAPK (64.7%), PI3K (66.7%), and SMAD (53.3%) signaling, and 17.6 to 40% of STAT-related pathways were independent from the other mechanisms (Fig. 1F).

### Mechanical pain sensitivity

#### Pressure application measurement (PAM) at the knee

Compared to baseline (pre-surgery) measures, the PAM threshold decreased (more pain sensitivity) at 4-weeks (β coefficient [95% confidence interval]) (−70.57g [−102.37, −38.77]), increased at 8-weeks (36.42g [4.62, 68.22]), and returned to baseline at 12-weeks (0.37g [−31.43, 32.17]) after OA induction surgery followed by control liposome (Veh-lip) injections (Fig. 2A, Sup. 6A).

**Figure 2.**
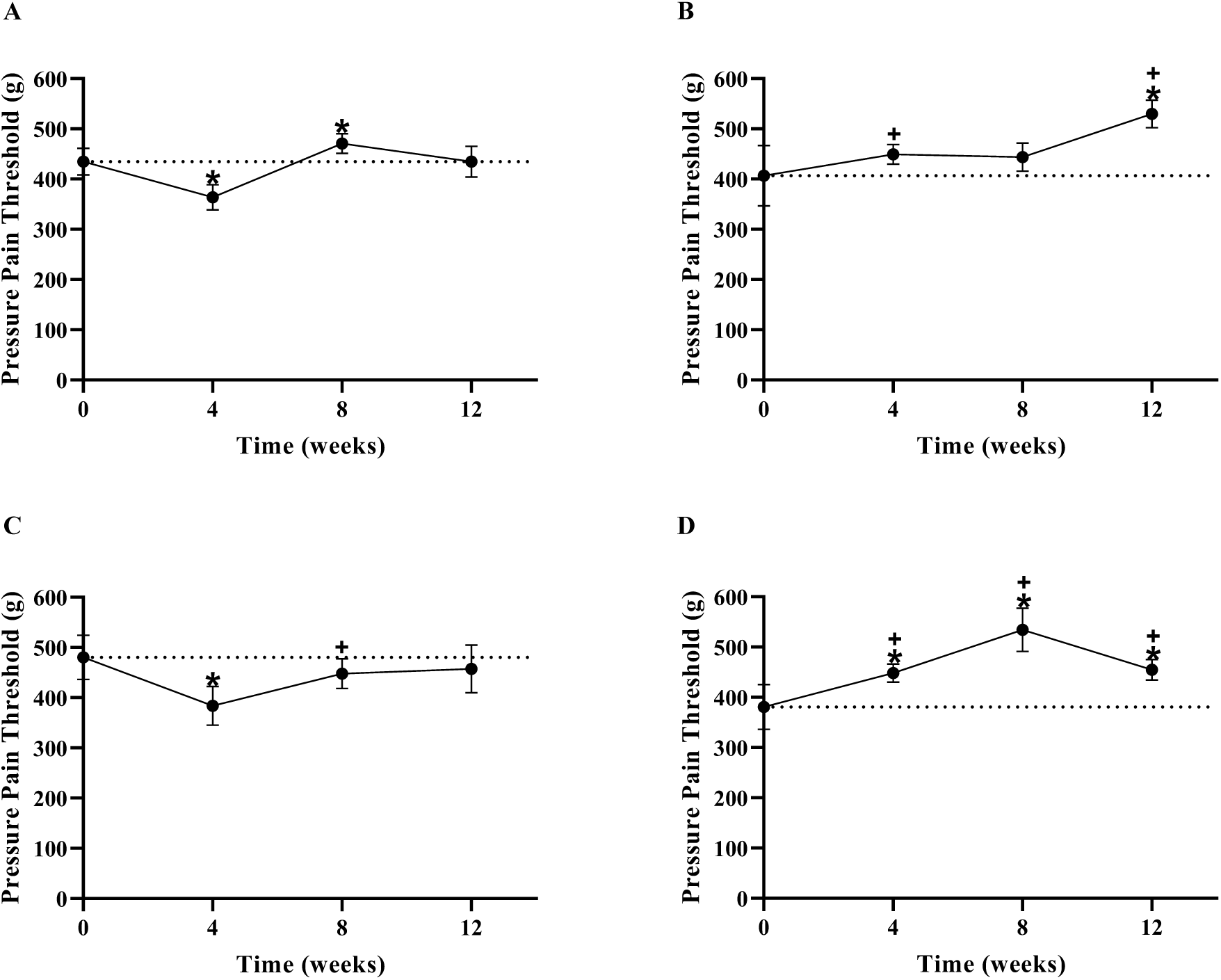
Knee pressure pain threshold as measured by pressure application measurement. Pressure pain threshold of Veh-lip (control) **(A)**, Clod-lip **(B)**, STAT1i-lip **(C)**, and STAT6i-lip **(D)** treatment at pre-surgical baseline (0), and 4-, 8- and 12-weeks post OA induction. The y-axis represents force (grams, g) applied to the knee before withdrawal and the x-axis time in weeks. Mean with 95% confidence intervals are displayed. Significant differences versus baseline (*) and versus control (Veh-lip) (+) are shown (P <0.05).

Macrophage depletion using clodronate-laden liposomes (Clod-lip) prevented a decrease in the PAM threshold at 4-weeks (42.18g [−3.27, 87.63]), and increased (improved) the threshold at 12-weeks (123.32g [77.87, 168.77] compared to baseline (Fig. 2B, Sup. 6B). Compared to Veh-lip controls, Clod-lip increased the PAM threshold at 4- (112.75g [50.60, 174.90] and 12-weeks (122.95g [60.80, 185.10]) (Fig. 2B, Sup. 7).

STAT1-inhibitor liposomes (STAT1i-lip) did not prevent a decrease in the PAM threshold at 4-weeks (−96.64g [−146.19, −47.09]), and prevented an increase in the threshold at 8- (−32.44g [−81.99, 17.11]) compared to baseline (Fig. 2C, Sup. 6C). Compared to Veh-lip controls, STAT1i-lip decreased the PAM threshold at 8-weeks (−68.86g [−131.01, −6.71]) (Fig. 2C, Sup. 7).

STAT6-inhibitor liposomes (STAT6i-lip) increased the PAM threshold at 4- (67.86g [6.38, 109.34]), 8- (153.85g [112.37, 195.33]), and 12-weeks (73.97g [32.49, 115.45]) compared to baseline (Fig. 2D, Sup 6D). Compared to Veh-lip controls, STAT6i-lip increased the withdrawal threshold at 4- (138.43g [76.28, 200.58]), 8- (117.43g [55.28, 179.58]), and 12-weeks (73.60g [11.45, 135.75]) (Fig. 2D, Sup. 7).

#### Hindpaw withdrawal threshold

Compared to baseline (pre-surgery) measures, the hindpaw withdrawal threshold decreased at 4- (−9.18g [−17.18, −1.18]), 8- (−8.23g [−16.23, - 0.022]), and 12-weeks (−12.93g [−20.93, −4.93]) after OA induction surgery followed by Veh-lip injections (Fig. 3A, Sup. 8A).

**Figure 3.**
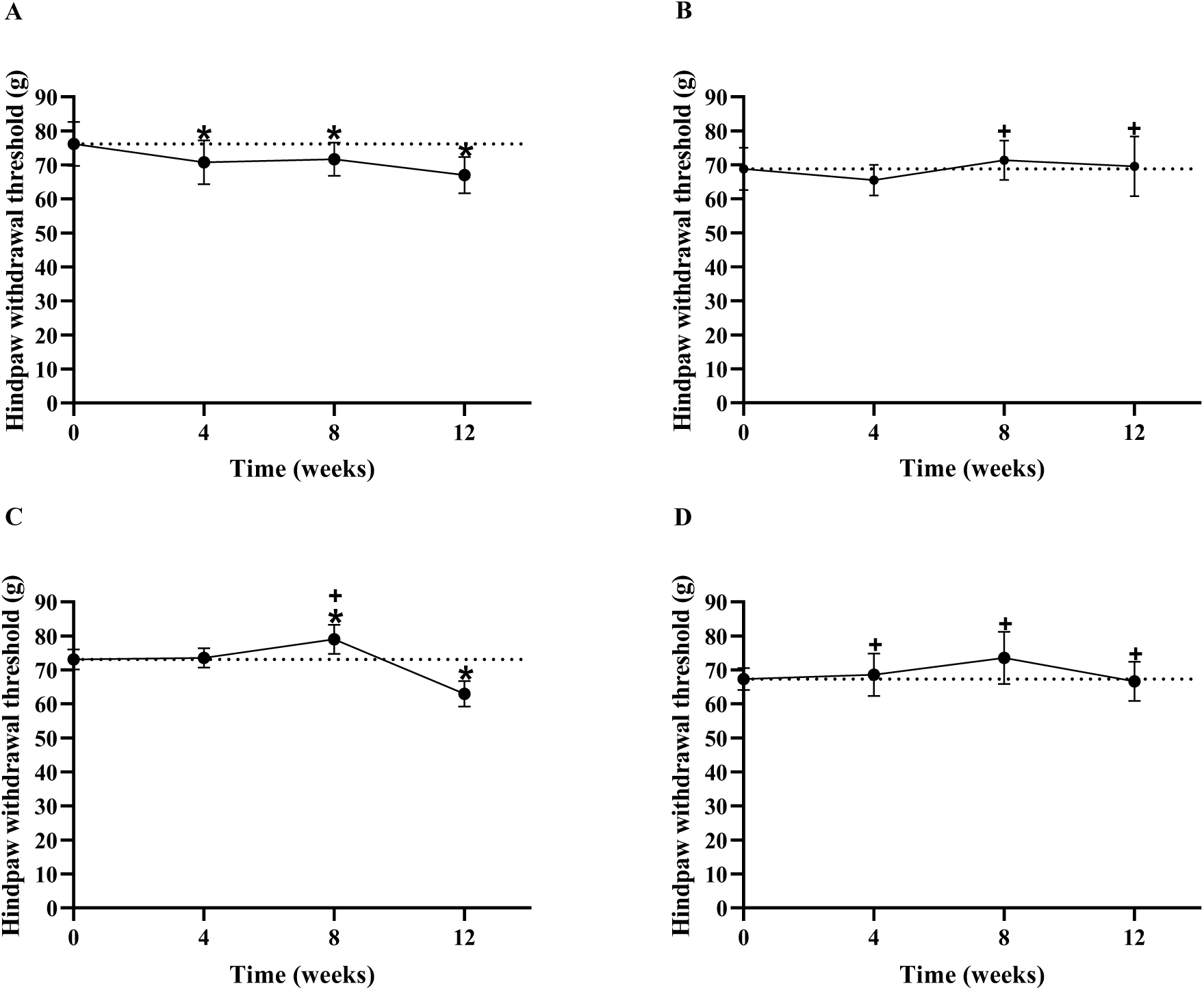
Ipsilateral hindpaw mechanical sensitivity measured by electronic von Frey withdrawal threshold. Hindpaw withdrawal threshold of Veh-lip (control) **(A)**, Clod-lip **(B)**, STAT1i-lip **(C)**, and STAT6i-lip **(D)** treatment at pre-surgical baseline (0), and 4-, 8- and 12-weeks post OA induction. The y-axis represents force (grams, g) applied to the hindpaw before withdrawal and the x-axis time in weeks. Mean with 95% confidence intervals are displayed. Significant differences versus baseline (*) and versus control (Veh-lip) (+) are shown (P <0.05).

Clod-lip injections prevented reductions in hindpaw withdrawal threshold at all time points compared to baseline. Compared to Veh-lip controls, Clod-lip injections increased the withdrawal threshold at 8- (10.83g [0.8, 20.85]), and 12-weeks (13.75g [3.73, 23.79]) (Fig. 3B, Sup. 9).

STAT1i-lip prevented a decrease in hindpaw withdrawal threshold at 4 weeks, increased the threshold at 8-weeks (5.89g [1.71, 10.07]), but did not prevent a decrease in the withdrawal threshold at 12 weeks (−10.16g [−14.34, −5.98]) compared to baseline (Fig. 3C, Sup. 8C). Compared to Veh-lip controls, STAT1i-lip increased withdrawal threshold only at 8 weeks (14.12g [4.09, 24.14]).

STAT6i-lip injections prevented decreases in withdrawal threshold at all time points compared to baseline. Compared to Veh-lip controls, STAT6i-lip increased withdrawal thresholds at 4- (10.45g [0.42, 20.47]), 8- (14.43g [4.40, 24.46]), and 12-weeks (12.26g [2.23, 22.28]) (Fig. 3D, Sup.9).

### Histopathology

#### Synovial histopathology

Representative synovial histopathology micrographs are shown in Fig. 4A. Compared to Veh-lip controls, macrophage depletion with Clod-lip decreased synovial lining thickness, sub-intimal infiltration, and fibrin deposition at 4 weeks (Fig. 4B), but increased synovial vascularization, perivascular edema, and fibrosis at 12 weeks post-OA induction (Fig. 4C). No changes in synovial histopathology measures were observed with STAT1 inhibition at either time point. STAT6 inhibition (STAT6i-lip) decreased synovial lining thickness at 4 weeks post-OA induction.

**Figure 4.**
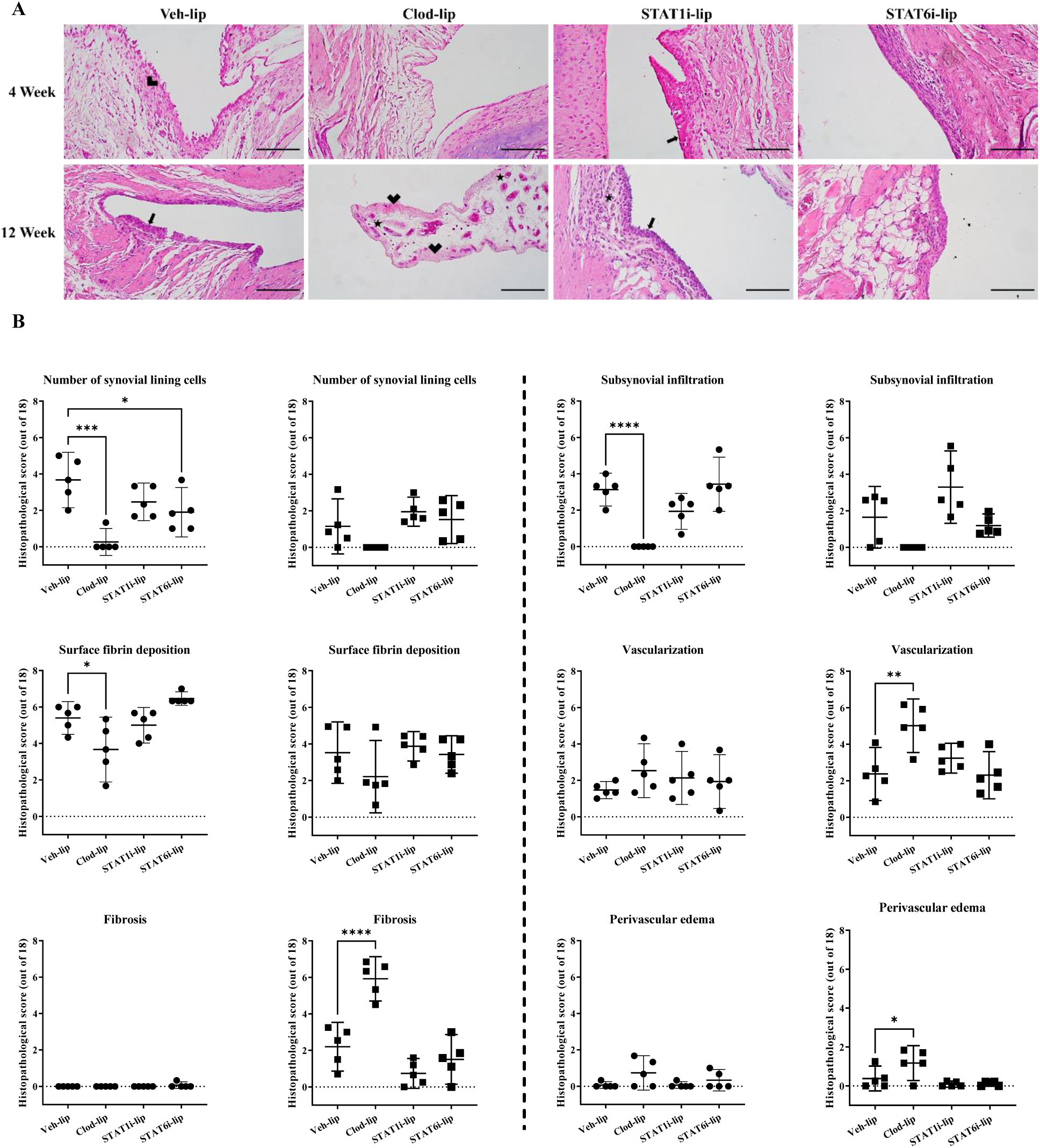
Synovial histopathology and synovitis grading during experimental OA development. Representative images of H&E-stained synovium at 4- and 12-weeks post surgery **(A)**. The scale bar represents 100µm, the thick arrow; synovial hyperplasia, thin arrow; sub synovial infiltrate, star; angiogenesis, and arrowhead; synovial fibrosis. Individual measures of synovial histopathology at 4-weeks (●) and 12-weeks (▪) post surgery with comparisons to Veh-lip. The y-axis shows the total histopathological score (out of 18) and the x-axis the treatment group. Mean with 95% confidence intervals are displayed, **** p<0.0001, *** p<0.001, ** p<0.01, * p<0.05.

#### Cartilage histopathology

Representative cartilage histopathology micrographs are shown in Fig. 5A. Proteoglycan loss, fissuring, and partial thickness cartilage erosion was focused in the central regions of articular surfaces at levels consistent with previous literature in this experimental model of OA, with progression in severity from 4- to 12-week time points (*15*). There were no changes in cartilage degeneration scores with any treatment, although a trend toward a decrease in total articular degeneration score was observed at 12 weeks post-OA induction in the STAT1i-lip treatment group in particular (Fig. 5C).

**Figure 5.**
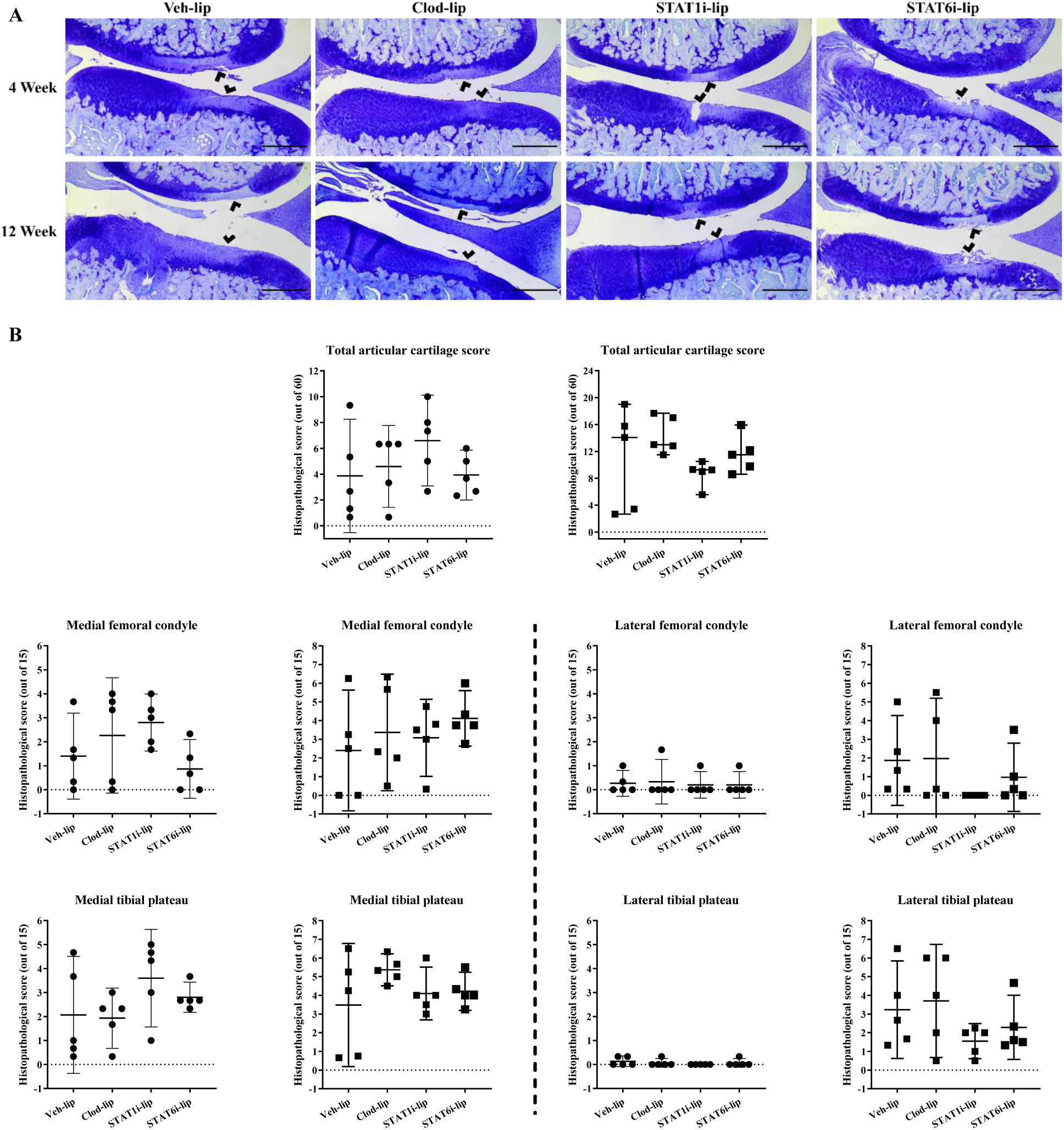
Cartilage histopathology and degeneration grading during experimental OA development. Representative images of toluidine blue stained synovium at 4- and 12-weeks post surgery **(A)**. The scale bar represents 500µm, and the arrowhead areas of cartilage degeneration. Measures of cartilage degeneration in the medial and lateral tibial plateau and femoral condyles at 4-weeks (●) and 12-weeks (▪) post surgery. The y-axis shows the total histopathological score (out of 60) or individual histopathological score (out of 15) and the x-axis the treatment group. Mean with 95% confidence intervals are displayed.

### Immunofluorescence detection of CD68+ macrophages in synovial tissue

Representative images of CD68+ macrophages in synovial tissue are shown in Fig. 6A. Intimal macrophage density remained constant whereas subintimal macrophage density increased at 12 weeks in the Veh-lip condition (Fig 6D, E). Clod-lip treatment reduced intimal macrophage density at 4-weeks post surgery vs Veh-lip (cells/mm^2^ [minimum, maximum]) (1.7 [0, 6.8] and 10.5 [2.6, 16.6] respectively; p=0.03) (Fig 6B). STAT1i-lip treatment caused no detectable effect on macrophage density in the intima. STAT6i-lip increased macrophages in the intima compared to Veh-lip (19.9 [4.5, 26.9] and 10.5 [2.6, 16.6] respectively; p=0.03). STAT6i-lip also increased subintimal macrophages 4-weeks after OA induction vs Veh-lip (3.4 [1.0, 8.4] and 0.3 [0, 1.1] respectively, p=0.002). No other treatment caused a detectable effect on subintimal macrophage density.

**Figure 6.**
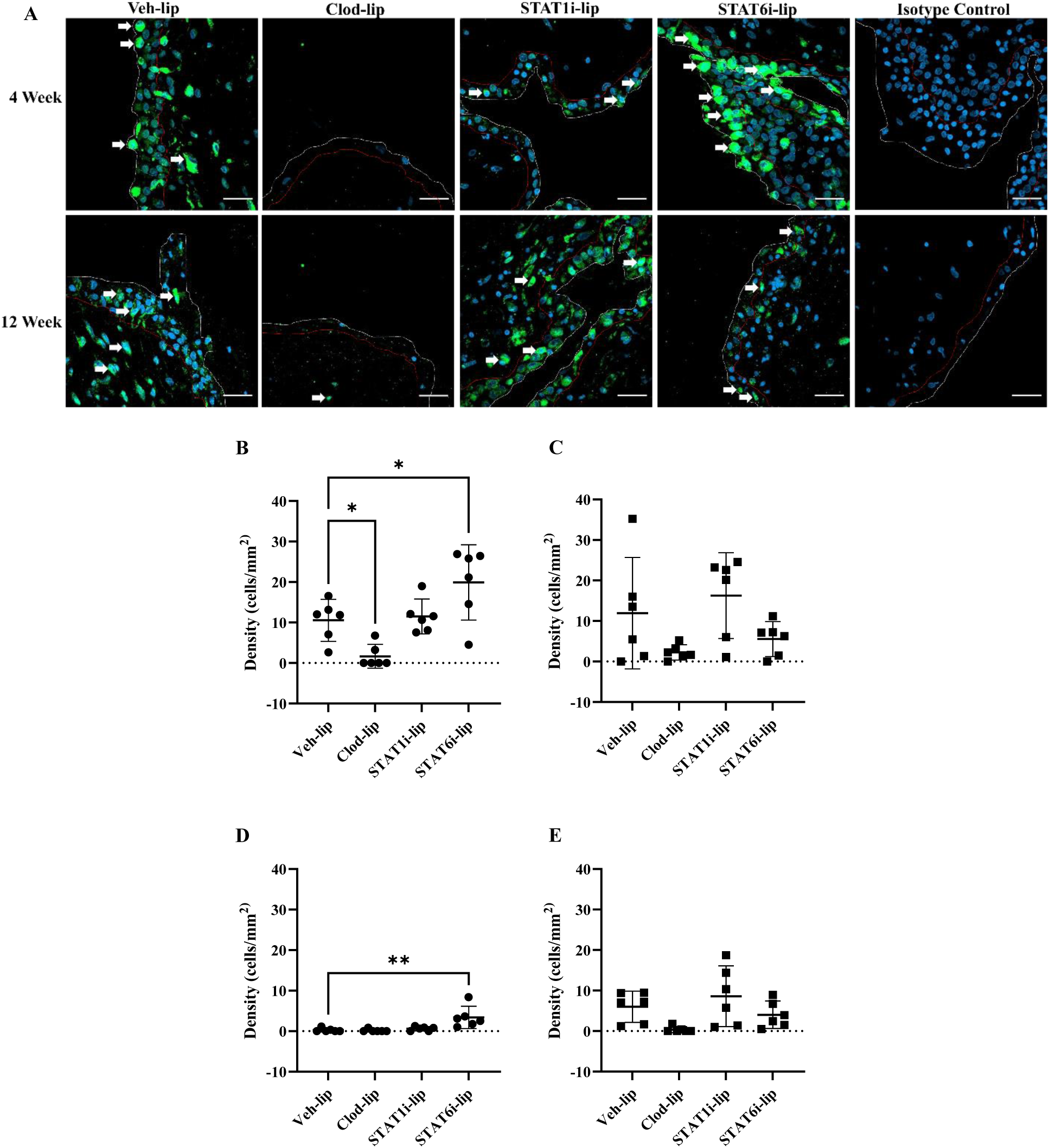
Macrophage quantification in the synovial intima and sub-intima. Representative confocal images of anti-CD68 stained (green) macrophages at 4- and 12-weeks post surgery **(A)**. The scale bar represents 25µm. The white line separates the edge of the synovium and joint space and red line the interface between intima and subintima. White arrows highlight CD68 positive cells. Cell density in the intima 4- (●) **(B)** and 12-weeks (▪) **(C)** and subintima 4- (●) **(D)** and 12-weeks (▪) **(E)** post surgery with comparisons to Veh-lip. Mean with 95% confidence intervals are displayed, ** p<0.01, * p<0.05.

### Sulfated glycosaminoglycan production and gene expression in articular chondrocytes co-cultured with synovial tissue

Sulphated glycosaminoglycan (sGAG) production by healthy articular chondrocytes increased in co-culture with STAT1i-lip-treated synovial tissue (compared to Veh-lip) collected from the 4-week time point (µg/mL [95% CI]) (6.43 [7.69, 5.16] and 4.72 [5.84, 3.57], respectively; p=0.03) (Fig. 7A). Although a trend toward increased sGAG secretion was observed in co-culture with STAT6i-lip-treated synovium from the same time point, the difference was smaller and lacked precision to detect a significant difference (Fig. 7A). No differences in sGAG secretion were seen when chondrocytes were co-cultured with synovial tissues from the 12-week time point.

**Figure 7.**
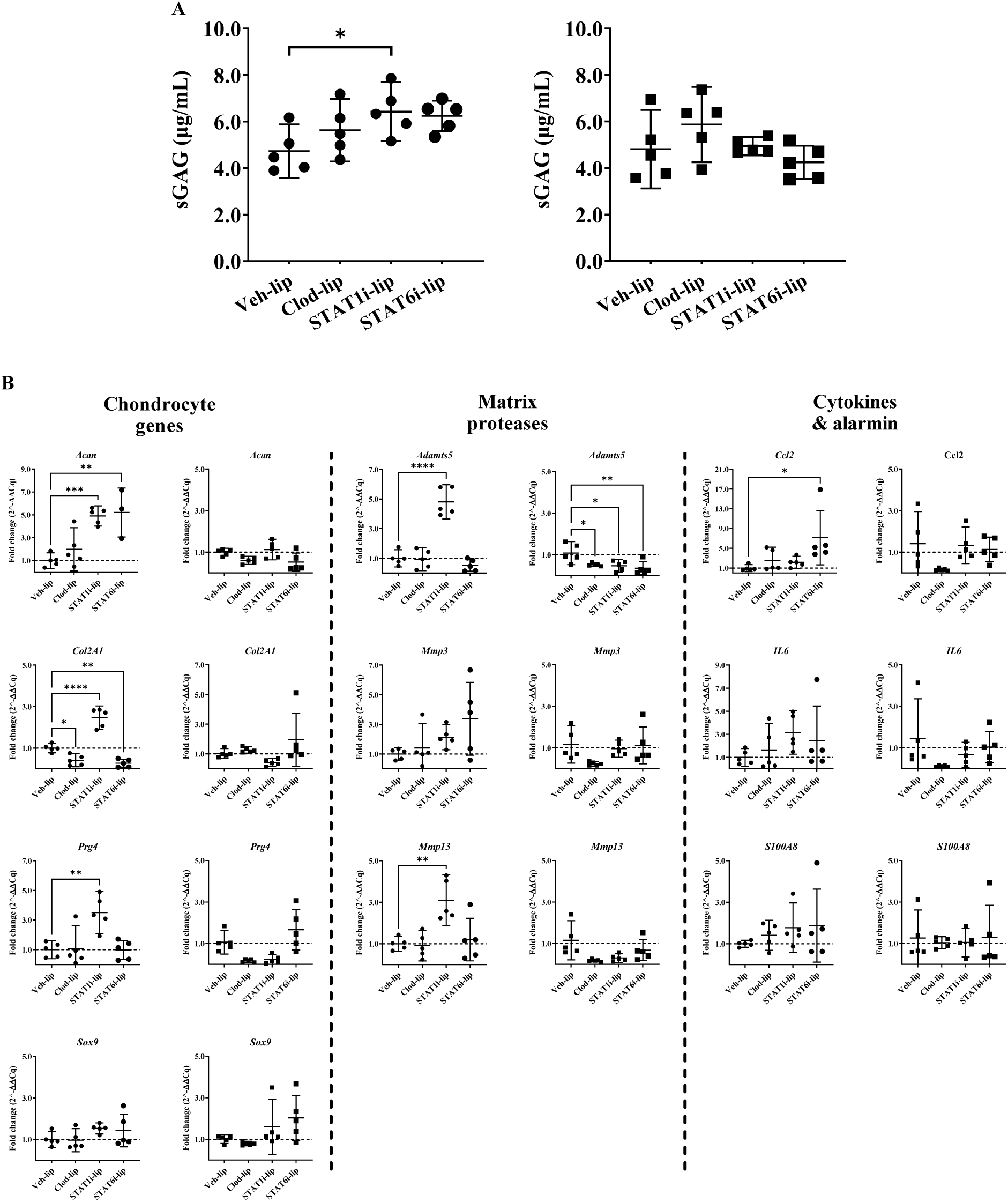
Effects of macrophage-targeted treatment on articular chondrocyte sGAG secretion and gene expression in co-culture with synovial tissue explants. sGAG production by chondrocytes after co-culture with synovial tissue explants from experimental OA knees treated with liposomal treatments for 4- (●) and 12-weeks (▪) **(A)**. The Y-axis represents concentration (µg/mL) of sGAGs in media collected after 24 hours of co-culture, * p=0.03. Gene expression of naïve chondrocytes co-cultured with synovial explants from 4- (●) and 12-week (▪) animals **(B)**. The y-axis represents fold change (2^-ΔΔCq) and the x-axis treatment group. Mean with 95% confidence intervals are displayed, **** p<0.0001, *** p<0.001, ** p<0.01, * p<0.05.

Most chondrocyte gene expression changes occurred in response to co-culture with synovial tissues collected from the 4-week time point. Compared to Veh-lip controls, Clod-lip synovium decreased *Col2a1* expression (fold change [upper and lower limit]) (0.4 [0.7, 0.1]; p=0.02) in chondrocytes, but no changes were observed in other ECM, protease, or cytokine/chemokine genes (Fig. 7C). STAT1i-lip synovium increased *Acan* (4.9 [5.8, 4.0]), *Col2a1* (2.5 [3, 1.9]), *Prg4* (3.5 [4.9, 2.1]), *Adamts5* (4.8 [6.0, 3.7]), and *Mmp13* (3.1 [4.3, 1.9]) expression (Fig. 7C). STAT6i-lip synovium increased A*can* (5.2 [10.5, 0]) and *Ccl2* (7.2 [14.0, 0.3]), and decreased *Col2a1* (0.3 [0.5, 0.03]) expression (Fig. 7C). Using synovial tissue from the 12-week time point, Clod-lip, STAT1i-lip, and STAT6i-lip treatment decreased *Adamts5* expression (0.5 [0.8, 0.2], 0.4 [0.7, 0], and 0.5 [0.6, 0.4] respectively) in chondrocytes compared to Veh-lip controls (Fig. 7D).

## Discussion

Chronic knee pain is the most commonly reported symptom by patients suffering from OA, but existing treatments are limited by side effects and safety issues (*16–18*). Synovitis is strongly associated with OA pain experiences (*3, 19*). Resident and recruited synovial macrophages are thought to play a role in nociception, but the mechanisms are not well understood. We used transcriptomics to identify pathways associated with synovial macrophage activation during experimental OA development, when nociception emerges. We found that STAT signaling is featured prominently during macrophage activation in early-stage experimental OA, and substantially overlaps with other major intra-cellular signaling pathways. Since the effects of STAT1 and STAT6 are mostly exclusive, we tested the role of STAT1- and STAT6-mediated macrophage activation in OA-related nociception, and further compared these with macrophage depletion. Macrophage depletion protected against the development of mechanical pain sensitivity, indicating that synovial macrophages mediate nociception during early-stage experimental OA development and progression. However, these benefits came at the cost of increased joint tissue damage including increased synovial fibrosis, vascularization, and perivascular edema, which are associated with increased radiographic OA severity in patients (*20*). These results show that macrophages are important mediators of mechanical pain sensitivity but are also required to maintain joint homeostasis. More selective treatments are therefore needed to reduce pain without causing tissue damage.

Strikingly, STAT6i in synovial macrophages raised (improved) the threshold for mechanical sensitivity above baseline and control levels at the affected knee and distal hindpaw at all time points, providing robust protection against mechanical pain sensitivity. Unlike macrophage depletion, we observed no worsening of synovial or cartilage histopathology with STAT6i, which also protected against synovial lining hyperplasia. On the other hand, STAT1i transiently lowered (worsened) knee withdrawal thresholds and increased hindpaw withdrawal thresholds. No clear impact of STAT1i on synovial or cartilage histopathology was seen, although trends toward a small chondroprotective effect were suggested at the 12-week endpoint. These results suggest that differential roles are played by STAT1 and STAT6 signaling with respect to macrophage activation, nociception, and joint tissue pathology in OA.

Macrophages are one of the most important synovial cell types in OA. Although studies have identified altered macrophage-related gene expression from whole synovial tissue (*21*), our study is among the first to explore differential gene expression directly in the synovial macrophage compartment during experimental knee OA development. In line with longitudinal gene expression studies in mouse models (*22*), synovial macrophage gene expression followed a phasic pattern. Early-stage OA macrophage gene expression reflected changes in cellular metabolism, activation, motility, inflammation, and Wnt signaling, which transitioned to mainly extracellular matrix remodeling after 12 weeks of OA development. Similar to McCoy *et al.* who identified increased cellular metabolism in synovial tissue transcriptomic analyses in an equine model of OA (*23*), our gene set enrichment analyses suggested a major role for STAT signaling in mediating cell stress, metabolism, and angiogenesis, with major overlaps between STAT and MAPK, PI3K, and SMAD signaling. STAT signalling may be a critical mediator of macrophage activation during early OA development.

A role for STAT signaling in OA cartilage has also been described. *Latourte et al.* demonstrated structural protection against joint destabilization-induced experimental knee OA in the mouse using prophylactic systemic inhibition of STAT3 (*24*). However, the effects of STAT inhibition on OA-related pain are not well-understood and the role of STATs in regulating macrophage activation is context-dependent (*25*). Given the complementary roles of STAT1 and STAT6 signaling in regulating macrophage activation, we chose highly-selective inhibitors of STAT1 (fludarabine) and STAT6 (AS1517499) for targeted delivery to synovial macrophages via phagocytosis of drug-loaded liposomes after intra-articular injection.

The potential for dual roles played by macrophages in mediating OA outcomes has long been suspected, partly based on the well-described model of pro- (M1) and anti-inflammatory (M2) polarization. However, our data underscore the hazards of relying on the M1/M2 paradigm, or assuming that all inflammation is bad/anti-inflammatory processes are good. For example, interleukin-4 (IL-4) is an anti-inflammatory cytokine and generally regarded as analgesic (*26*). IL-4 knock-out mice display increased hindpaw mechanical hypersensitivity and STAT6 is a key mediator of IL-4 receptor signaling (*25, 27*). In contrast, we found that STAT6i robustly protected against the development of both distal and local pain, suggesting that STAT6 activation likely has alternative functions in OA synovial macrophages. Interestingly, Haraden *et al.* reported that OA disease severity was correlated to synovial fluid levels of the M2-marker CD163 (*28*), which aligns with our results and suggests that alternatively-activated macrophages may contribute to nociception. Additionally, atopic diseases characterised by STAT6 activation increase OA incidence (*29*), highlighting its importance in OA pathogenesis. Arguing against a major role for pro-inflammatory macrophages as key mediators of nociception in OA, an early, brief benefit on distal pain sensitivity along with worsening of local knee pain was observed in STAT1i. STAT1 signaling may therefore play a smaller role in mediating OA-related nociception in the early stages after injury.

Analgesic treatments are frequently associated with off-target effects. For example, NGF inhibition caused a small but important increased risk of rapidly progressive OA (*1*). To ensure that macrophage-targeted treatments did not worsen joint tissue outcomes, we explored joint histopathology and crosstalk between treated synovial tissue and healthy primary articular chondrocytes in our established *ex-vivo* joint co-culture system (*15*). Importantly, no treatment led to any increase in articular cartilage damage at individual articular surfaces or overall. There were trends toward reduced cartilage damage with STAT1i and STAT6i, but we lacked statistical power to detect a small protective effect. Thus, we cannot entirely exclude the potential for a protective effect on joint damage, as Sun *et al.* reported marked reduction in cartilage degeneration after depleting macrophages in obese mice prior to, and 1 week after, OA induction (*30*). The prophylactic strategy in that study may also have led to an increased effect.

We previously reported that synovial tissue from early-stage experimental knee OA stimulates a transient anabolic response in healthy articular chondrocytes (*15*). In similar crosstalk analyses in this study, macrophage-depleted synovium from early-stage OA (4 weeks) decreased *Col2a1* expression in healthy articular chondrocytes, suggesting that synovial macrophages partially stimulate anabolic chondrocyte responses. In contrast, STAT1i caused increased sGAG secretion and expression of matrix (*Acan*, *Col2a1*, and *Prg4*) and protease genes (*Adamts5* and *Mmp13*). This suggests that STAT1-mediated macrophage activation in early-stage OA may suppress anabolic and matrix turnover processes in chondrocytes. STAT6i in synovial macrophages led to increased *Acan* and *Ccl2* gene expression and decreased *Col2a1* expression. Overall, our data suggest that STAT1-mediated macrophage activation inhibits anabolic responses in chondrocytes, whereas STAT6-mediated macrophage activation may be more important for nociception with minimal effects on chondrocyte anabolism.

Beyond cartilage damage, synovial macrophage depletion caused worse synovial vascularization, fibrosis, and perivascular edema at 12 weeks post-OA induction, which was not seen with STAT1i or STAT6i. Other studies have found that transient and/or prophylactic depletion of macrophages resulted in worse synovitis, joint damage, and infiltration of T lymphocytes (*31, 32*). In those studies, depletion was performed at a single timepoint, whereas we used repeat dosing every 2 weeks to sustain macrophage depletion/suppression. Together with our findings, a clear picture is emerging that macrophages are essential for maintaining joint homeostasis, while simultaneously playing pathological roles in nociception and suppressing anabolic processes through complementary mechanisms. Rather than depleting macrophages, our study suggests that selectively inhibiting STAT-mediated macrophage activation may be an effective alternative. Further studies are warranted, including the potential for a synergistic effect of inhibiting both STAT1 and STAT6 concurrently.

Our study has limitations. Our surgically induced joint destabilization model of OA does not address other OA risk factors such as age, obesity, or female sex, and the effects of STATi should be confirmed in those settings. Other pain-related behavioural tests may have revealed different outcomes, although our study focused on two of the most commonly used methods to assess mechanical pain sensitivity at the knee and hindpaw. We cannot rule out a role played by other phagocytes such as dendritic cells, mast cells, or neutrophils (rarely seen in OA), which may also have been targeted by liposomes. However, synovial macrophages are the dominant immune cells in healthy joints and are heavily recruited to the synovium of OA joints. Further studies will be required to determine whether different roles are played by tissue resident versus recruited (bone marrow-derived) synovial macrophages.

Our findings demonstrate that synovial macrophages play a major role in mediating nociception related to mechanical pain sensitivity in experimental knee OA. Although depleting macrophages improved pain sensitivity, this came at the cost of increased joint tissue damage. Macrophages therefore likely play dual roles in pain and joint organ homeostasis, suggesting that macrophage “tuning” may be more successful than depletion. Selective targeting of macrophages with STAT inhibitors is a novel candidate strategy for pain modification in OA. STAT6i confers substantial protection against mechanical pain sensitivity without aggravating synovial histopathology, whereas STAT1i may improve cartilage anabolic responses if used very early after acute joint injury. Further studies are warranted to assess the effects of this treatment approach in other OA contexts.

## Methods

### Study design

To investigate the molecular mechanisms associated with synovial macrophage activation during early-stage experimental knee OA development and progression, we isolated synovial macrophages from the knee joints of rats at 4 and 12 weeks after surgical induction of knee OA via joint destabilization. Synovial macrophages were dissociated from synovial tissues and sorted out of single cell suspensions prior to bulk RNA isolation and sequencing. Comparisons at each time point were made to synovial macrophages isolated from a similar group of animals that underwent a sham surgery via knee joint arthrotomy without surgical joint destabilization. To test the effects of macrophage depletion, STAT1, or STAT6 inhibition in synovial macrophages during OA development and progression, we used drug-laden liposomes to selectively target these inhibitors to synovial macrophages via repeated intra-articular injections into the knees of a separate group of rats that underwent surgical induction of knee OA for up to 12 weeks. Primary outcomes included mechanical pain sensitivity measures at the knee and hindpaw. We assessed secondary outcomes including knee joint cartilage and synovial histopathology, macrophage infiltration, and crosstalk between synovium and articular chondrocytes in an ex vivo co-culture system. All experiments and replicates were performed with age-matched animals. All animals and samples were used for analysis. Endpoints were chosen based on previous research identifying OA disease phenotypes in this post traumatic rat model (*15*).

### Rat model of experimental knee OA

Sprague Dawley rats (Charles River Laboratory, Quebec, Canada, strain code 400) were housed and handled in the Animal Care and Veterinary Services conventional housing facility at Western University in accordance with the guidelines of the Canadian Council on Animal Care. The animal use protocol was approved by the Western University Animal Care and Use Committee (AUP2017-042). Twelve-week-old male rats underwent anterior cruciate ligament transection and destabilization of the medial meniscus surgery (OA), or a sham surgery (control) as previously described (*15*). Briefly, a medial arthrotomy of the right knee was performed followed by transection of the anterior cruciate ligament and the anterior medial meniscotibial ligament to induce joint destabilization (OA), or medial arthrotomy alone (sham surgery).

### Synovial cell and RNA isolation

Synovial tissue was dissected away from skin, tendon, muscle, fat pad, and menisci from the entire knee as previously described (*15*). Synovial tissues from 3 or 4 animals were pooled per replicate, to provide n=5 replicates per condition at 4 and 12 weeks after joint surgery. After rinsing in PBS, synovial tissue was minced in Roswell Park Memorial Institute (RMPI) 1640 media (Fisher Scientific, 11-875-093) containing 10% fetal bovine serum (FBS) (Fisher Scientific, 12483-020), 1% penicillin-streptomycin (Fisher Scientific, 15140-122), 940ug/mL of Liberase TM (Sigma-Aldrich, 5401127001) and 940ug/mL DNase I (Sigma-Aldrich, DN25). Cell dissociation reactions were incubated a 37° Celsius for 30 minutes with gentle rotation before 100mM EDTA was added to stop the reaction. Each reaction was then passed through a 100um cell strainer and centrifuged at 300g for 10 minutes. The synovial cell pellet was resuspended in Dulbecco’s Modified Eagles Medium (fisher Scientific, 11-885-084) containing 10% FBS, 1% penicillin-streptomycin before an aliquot was removed and diluted 1:1 with trypan blue to assess cell number and viability. CD11b/c positive cells (monocyte/macrophage) were sorted using magnetic-activated cell sorting (MACS) separation columns (Miltenyi Biotec, 130-105-634) and immediately lysed with TRIzol (Fischer Scientific, 15-596-018) for RNA isolation using the Direct-zol RNA MicroPrep Kit (Zymo Research, R2062) per the manufacturer’s protocol. RNA was analysed using a Qubit 1.0 fluorimeter (Thermo Fisher Scientific) using the Qubit RNA high sensitivity kit (Thermo Fisher Scientific, Q32852); RNA with RNA integrity numbers of 7.8-9.3 were used for sequencing.

### RNA-sequencing data analysis

RNA sequencing libraries were prepared with the Illumina stranded total RNA prep with ribo-zero plus kit (Illumina, 20040525). LabChip (Perkin Elmer, Caliper GX) was used for quality assessment and qPCR for quantitation of libraries. Libraries (100 base-pair, paired-end) were sequenced on 1 lane for 100 cycles from each end of the fragments on an Illumina NovaSeq 6000 using a NovaSeq 6000 S4 reagent kit v1.5 (Illumina, 20028313) at the McGill Genome Centre. The raw fastq files were downloaded from McGill Genome Centre. Fastp (version 0.20.0) was used to collect QC metrics of the raw reads (*33*).

Next, RNA sequences were aligned and sorted by coordinates, to the NCBI rat genome Rattus_Rnor6_V102, using STAR aligner (STAR-2.7.6a) (*34*). The removal of alignment duplicates was done with Sambamba (version 0.8.0) (*35*). Quantification of genes was performed using featureCounts (v2.0.0) (*36*). DESeq2 (v 1.24.0) was used to normalize feature counts and to test find the differentially expressed genes (*37*). The HGNC symbols were extracted and added to the DESeq2 results data frame using biomaRt (version 2.40.4) using the rat “rnorvegicus_gene_ensemb” BioMart version “Ensembl Release 102 (November 2020)” (*38, 39*). Differential gene expression for OA versus sham (control) included a log2-fold change ≥0.5 (absolute fold change ≥1) and an adjusted p value <0.05. Volcano plots were created using VolcaNoseR and bubbleplots in Rstudio (4.2.0) using ggplot2 (*40, 41*). Gene Set Enrichment Analysis (GSEA version 4.1.0) of normalized counts, was used to identify significantly enriched gene set/pathways between OA and sham, at each timepoint, using the Gene Ontology and Hallmark collections from the Molecular Signatures Database (MSigDB) (*42*). Significantly enriched pathways were considered as those with a false discovery rate of less than 0.05. Venny 2.1.0 was used to compare transcription factor involvement in enriched Hallmark pathways (*43*).

### Drug-loaded liposomes

Commercial liposomes containing phosphate buffered saline (Veh-lip; control) or 18.4 mM clodronate (Clod-lip) (Encapsula NanoSciences, CLD-8910), and liposomes prepared in-house containing fludarabine (STAT1i-lip) (Sigma-Aldrich, F9813) or AS1517499 (STAT6i-lip) (Sigma-Aldrich, SML1906) were used in this study. To match the STAT1i-lip and STAT6i-lip to the commercial liposomes, we prepared a spontaneously formed vesicle dispersion containing a 7:3 molar ratio of L-α-phosphatidylcholine (Avanti Polar Lipids, 840051P) and cholesterol (Sigma-Aldrich, C8667) in chloroform. The dispersion was subjected to rotary evaporation at 30 revolutions per minutes and 420 mbar in a 55°C water bath. Remaining traces of chloroform were removed under high vacuum. The phospholipid film was hydrated with a 2.5 mg/mL solution of fludarabine (STAT1i-lip) in PBS. AS1517499 was added prior to phospholipid film preparation due to its hydrophobic nature and STAT6i-lip encapsulated AS1517499 at 1 mg/mL. STAT1-lip and STAT6i-lip were extruded 6 times through 1.0 µm pore polypropylene filters to form a liposome suspension with an average diameter of 1.25 µm (0.7-2 µm) to match the commercial liposomes. Following extrusion, liposomes were purified using a 30,000 Dalton centrifugal size exclusion column (Sigma-Aldrich, UFC903008). Purified liposomes were resuspended at an equivalent lipid to buffer ratio prior to downstream analysis and injection. To validate shape and structure, early preparations of STAT1i-lip and STAT6i-lip were fixed in 1.5% paraformaldehyde and 1.5% glutaraldehyde in PBS and imaged on a Philips Electronics CM10 Transmission Microscope with a Hamamatsu ORCA 2MPx HRL Camera (Supplemental Fig. 1A). Dynamic light scattering (Malvern Panalytical, Zetasizer Nano ZS) was used to confirm size distribution in all batches of internally prepared liposomes and ensure a match to the commercially available preparations (Supplemental Fig. 1B).

### Liposome delivery

Liposomes were delivered via intraarticular injection. Rats were anaesthetized, the skin around the joint was prepared for injection using a sterile Hamilton syringe equipped with a 30-gauge needle to deliver 50 µL of liposomes. The needle was oriented to the central line of the joint and inserted 2-3 mm through the infra-patellar fat pad to reach the inter-condylar joint space. Injections were initiated 14 days after OA induction surgery, providing a total of 2 injections (day 14 and 21) in the group of animals collected at the 4-week endpoint, and 6 injections (day 14, 21, 28, 42, 56, and 70) in the 12-week endpoint group. We initiated our injections 14 days after OA induction to avoid interfering with the normal wound-healing process after surgical injury.

### Pressure Application Measurement (PAM)

Mechanical sensitivity at the OA knee was measured using pressure pain threshold longitudinally in all animals in the 12-week endpoint group (n=10 per condition) using a pressure application measurement (PAM) algometer (Bioseb, Boulogne, France). PAM was measured and reported in grams (g) at baseline, 4-, 8-, and 12-week time points by placing the algometer on the medial joint line of the knee and increasing pressure, approximately 200 grams per second, until a withdrawal or vocalization occurred. A 15-minute recovery period was given before a second measurement was taken, if needed.

### Electronic von Frey (eVF)

Using the same measurement approach as above, ipsilateral hindpaw mechanosensitivity was measured longitudinally in all 12-week endpoint rats using an electronic von Frey instrument (Bioseb). Rats were placed in an elevated metal-mesh floored Plexiglass chamber, allowing access to the plantar surface of the hindpaws. An acclimatization period was given until exploratory behaviour stopped (approximately 20 minutes). Measurements were taken by applying force to the plantar surface of the hindpaw with the eVF applicator tip, at a rate of 25 grams per second, until withdrawal occurred. Three measurements were taken with a 5-minute recovery period between each read. The average hindpaw withdrawal threshold was measured and reported in grams (g).

### Cartilage and synovial histopathology

Whole knees were harvested for histopathology analysis at 4- and 12-week endpoints from half of the rats included in the liposome treatment cohort (n=5 per liposome condition per timepoint). Tissue was fixed in neutral buffered 4% paraformaldehyde (Sigma-Aldrich, P6148-1KG) overnight at 4°C and decalcified in Decal Stat (StatLab, PC-1212-1) for 24 hours. Following this, joints were hemisected into anterior, containing the infrapatellar fat pad and para-articular tibial bone, and posterior portions, and further decalcified for 6 days in Formical-2000 (StatLab, PC-1314-32). Decalcified tissue was processed, embedded, and sectioned in the frontal plane. Twenty adjacent 5 µm sections were collected every 600 µm through the joint, with two adjacent sections prepared per slide. Adjacent sections every 600 µm were stained with toluidine blue (Fischer Scientific, AC348600250) or Harris Hematoxylin (Leica, 3801562) and Eosin Y (Fisher Scientific, E511) (H&E). Cartilage degeneration was graded on one toluidine blue stained slide, every 600 µm, using the OARSI Histopathology cartilage degeneration system described by Gerwin *et al.* (*44*). Briefly, the medial and lateral femoral condyles and tibial plateaus were split into inner, middle, and outer thirds. Each region was graded on a scale of 0 (no damage) to 5 (severe) and summed for a total score of 15 per articular facet. Synovial histopathology was graded on one H&E-stained slide, every 600 µm, using the six-component scoring system as described by Minten *et al.* (*20*).

Briefly, the medial and lateral parapatellar and tibiofemoral regions were each scored on a scale of 0 (none) to 3 (severe) for each of six components including synovial lining thickness (hyperplasia), synovial infiltration, fibrin deposition, vascularization, fibrosis, and perivascular edema, and each of which was summed for a total score of 18 per component. Slides were digitized on a Leica DM1000 with 40x magnification for toluidine blue stained slides and 200x magnification for H&E stained slides.

### Experimental joint tissue co-culture system

The remaining knees from the liposome treatment cohort were allocated to tissue co-culture testing (n=5 per condition). Whole knee synovial tissue was dissected under sterile condition and used in co-culture as previously described (*15*). Primary chondrocytes were harvested from the cartilage of a separate group of healthy (non-OA) rats. Cartilage was shaved from the femoral and tibial articular surfaces and digested in DMEM with collagenase II (Fisher Scientific, 17101015), DNase I, and mechanical force. Chondrocytes were expanded, pooled, and cryogenically frozen to reduce cell population variability in the co-culture system. P1 chondrocytes were thawed, seeded at 10% density, and allowed to reach confluency prior to serum-starving for 16 hours and assembly into co-cultures with liposome-treated synovial tissues. Whole synovium from a single knee was placed into a seated 12-well trans-well insert (Fisher Scientific, 353494) with serum free 1:1 DMEM/F12 with 1% penicillin-streptomycin in both chambers and equilibrated for 24 hours before transfer into co-culture. To assemble co-cultures, synovium and conditioned medium was transferred *en bloc* into co-culture for 24 hours with chondrocytes. For analysis, conditioned medium was collected to assess sulfated glycosaminoglycan production, synovium was fixed for immunofluorescent analysis, and chondrocytes were lysed with TRIzol for RNA purification and gene expression analysis.

### Sulfated glycosaminoglycan assay

Sulfated glycosaminoglycan (sGAG) content in conditioned medium was assessed as previously described (*15*). In that study, we confirmed that sGAG levels in this system reflect increased synthesis of GAG by chondrocytes by co-measurement of CS 846, a marker of newly-synthesized aggrecan. Conditioned medium or controls were diluted 1:2 with dimethyl-methylene blue (Sigma Aldrich, 341088) and absorbance at 595nm was measured on a spectrophotometer.

### Chondrocyte RNA isolation and gene expression analysis

Chondrocyte RNA purification and gene expression analysis was performed as previously described (*15*). TRIzol was used to lyse chondrocytes from co-culture experiments. Phase separation tubes (Fisher Scientific, A33248) and the RNAEasy Mini Kit (Qiagen, 74106) were used to purify RNA. iScript Reverse Transciption Supermix (Bio-Rad, 1708841) was used for reverse transcription, and quantitative real-time PCR was performed using SsoAdvance Universal SYBR Green Supermix (Bio-Rad, 1725274) with predesigned primers for *Acan, Col2A1, Prg4, Sox9, Adamts5, Mmp3, Mmp13, Ccl2, Il6, and S100A8* (Bio-Rad). Cq values were calculated on CFXMaestro 1.1 (Bio-Rad) and normalized to reference genes prior to analysis using the 2^(−ΔΔCq)^ method.

### Immunofluorescence

Synovial macrophages were assessed in dissected synovium from co-culture experiments (described above) that was fixed after in-vitro experimentation, processed, and embedded in Paraplast (Fisher Scientific, 23-021-750). Slides containing two 5 µm serial sections were prepared and baked at 60°C for 20 minutes prior to deparaffinization. Two slides per condition (n = 3 per liposome condition) was immersed in 70°C TRIS antigen retrieval buffer (10 mM TRIS, 1 mM EDTA, 10% glycerol, pH 9) and allowed to cool for 1.5 hours.

Permeabilization was with 0.2% Triton-x 100 (Sigma-Aldrich, 9002-93-1) in PBS and blocked with 5% bovine serum albumin (Sigma-Aldrich, A3294) and 0.1% Triton-x 100 in PBS. One section per slide was probed with rabbit anti-CD68 (Abcam, ab125212) and the other with rabbit polyclonal isotype control (Abcam, ab37415) at 0.005 mg/mL in blocking buffer overnight at 4°C. Goat anti-rabbit-Alexa-488 secondary antibody (Jackson ImmunoResearch, 111-545-144) at 0.0015 mg/mL was applied before washing and mounting with ProLong Gold Antifade Mountant with DAPI (Fischer Scientific, P36931). Slides were imaged on a Zeiss LSM 800 AiryScan confocal microscope using the 40X water immersion lens with configuration settings held constant across all samples. Two regions of interest were imaged per tissue section, providing at least 4 high power fields for counting the number of CD68+ cells in the synovial intima and sub-intima normalized to area.

### Statistical Analysis

A linear mixed effects model with alpha error at 0.05 determined that a sample size of 8 per condition would provide at least 80% power to detect a moderate difference in pain between groups. To account for additional degrees of freedom for treatments, we included a sample size of n=10 per condition for the primary analysis of mechanical pain sensitivity measures. To assess changes in mechanical sensitivity measures, individual linear mixed effects regression models were used for each treatment group (Veh-lip, Clod-lip, STAT1i-lip, STAT6i-lip) to determine changes in pain behaviours while accounting for the effect of time. A separate model was fit for each pain behaviour assessment (PAM and eVF). Restricted maximum likelihood estimation was used to account for missing data. In each model, time (measured in weeks post-model induction) was entered as a categorical predictor variable for fixed effects. Random intercepts were specified by rat ID. Assumptions for linear mixed models were tested; residuals were normally distributed and homoscedastic for both models. Likelihood ratio tests (LRTs) and Bayesian Information Criterion (BIC – lowest values preferred) were used to evaluate model fit and to select appropriate model covariance structure to control for correlation among repeated within-subject measures. Measures of association are reported as unstandardized beta (β) coefficients ± 95% confidence intervals (CIs) and standard error. In addition, post estimation pairwise comparisons of treatment to control (Veh-lip) were completed with Sidak correction. Cartilage and synovial histopathology, synovial macrophage density, and chondrocyte co-culture measures of sGAG and gene expression were analyzed with one-way ANOVA in GraphPad Prism (version 9.1.2) with Dunnett multiplicity adjustment. A p-value of ≤0.05 was considered significant.

## Supplemental material (list)

- Supplemental_figures_STATi_Manuscript.docx
- Supplemental_tables_1-5_STATi_Manuscript.docx
- Supplemental_tables_6-9_STATi_Manuscript.xlsx

## Acknowledgements

We thank the Biotron Integrated Microscopy Facility for help with transmitted electron microscopy imaging. We would also like to thank Joshua Jadischke for their assistance with liposomal extrusion.

## Contributions

Study conception and design. YLZ, KKP, ERG, and CTA. Acquisition of data. GB, YLZ, JK, KKP, AM. Analysis and interpretation of data. GB, YLZ, HTP, BF, LAW, ERG, CTA. All authors were involved in drafting the article or revising it critically for important intellectual content, approved the final version submitted, and take responsibility for the integrity of the data and the accuracy of the data analysis as per ICJME guidelines.

## Data

RNA sequencing data files were deposited in the Gene Expression Omnibus (GEO) at the NCBI (GSE216932). All other datasets used for the present study are available from the corresponding author upon reasonable request.

