## Supplemental figures for "Synovial macrophage activation mediates pain experiences in experimental knee osteoarthritis"

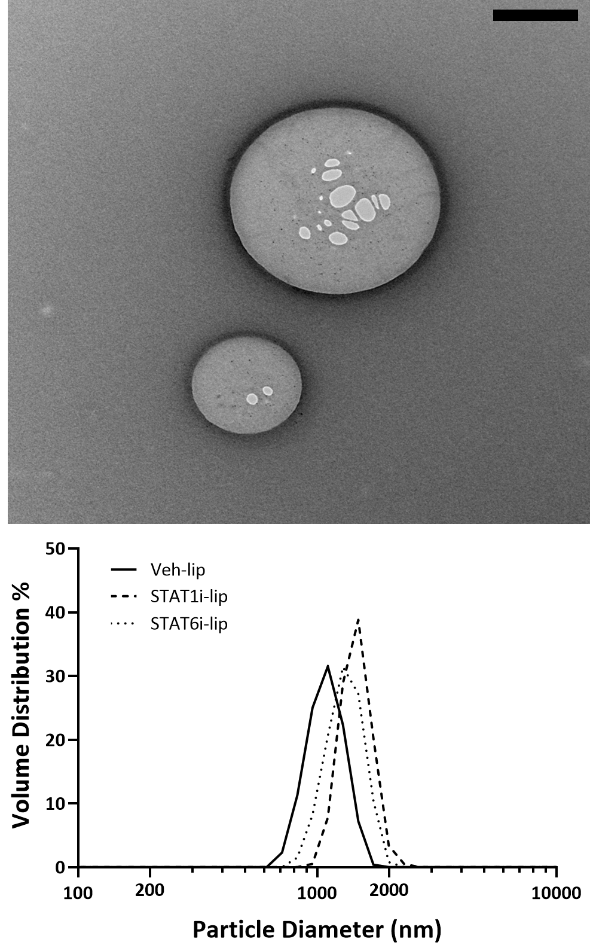

**Size distribution (%)**

**Particle diameter (nm)**

**B**

**A**

**Supplemental Fig 1. Liposome shape and size distribution.** Representative photomicrograph of prepared drug containing liposome **(A)**. Scale bar represents 0.5 µm. Representative dynamic light scattering results are displayed **(B)** with the y-axis showing volume distribution (% of total population) and x-axis the particle diameter.
