## Supplemental tables 6-9 for "Synovial macrophage activation mediates pain experiences in experimental knee osteoarthritis"

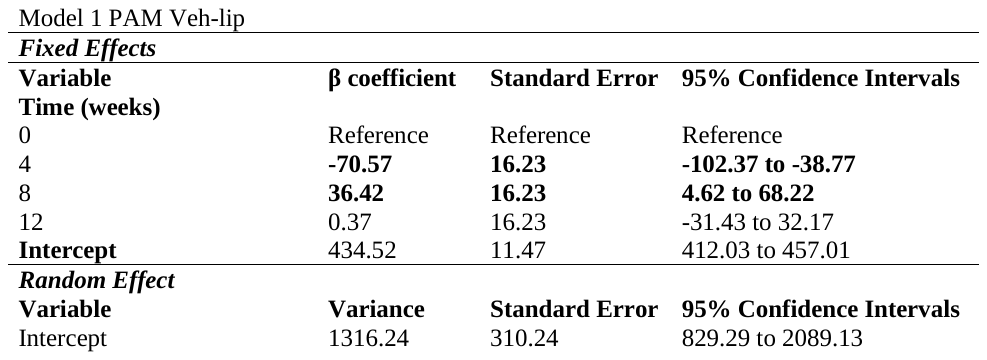

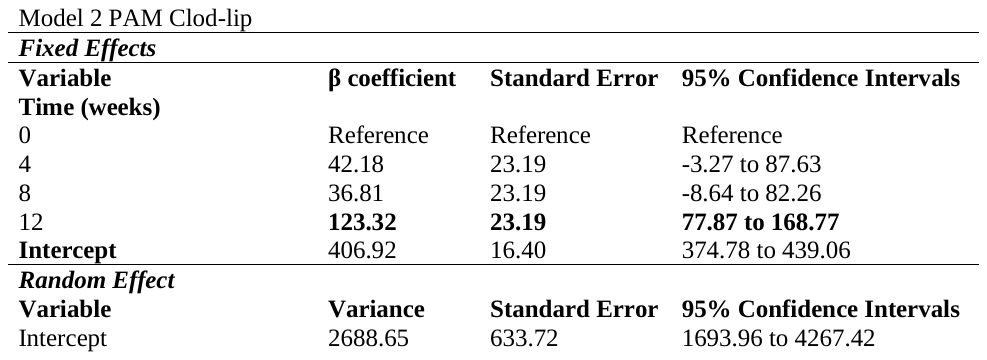

**C**

**A**

**B**

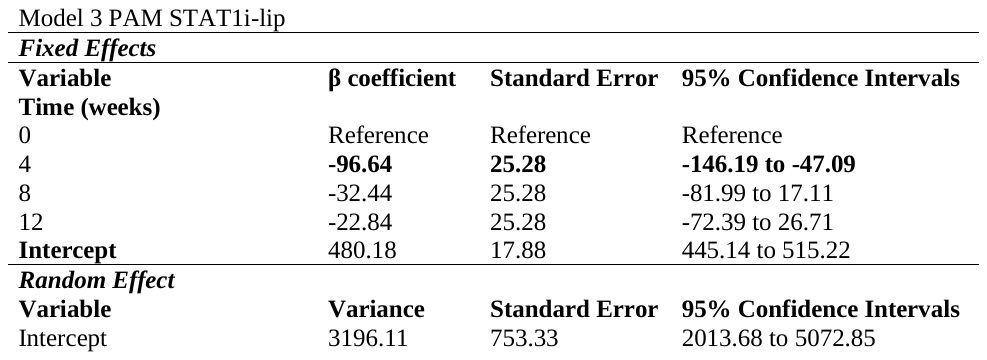

**D**

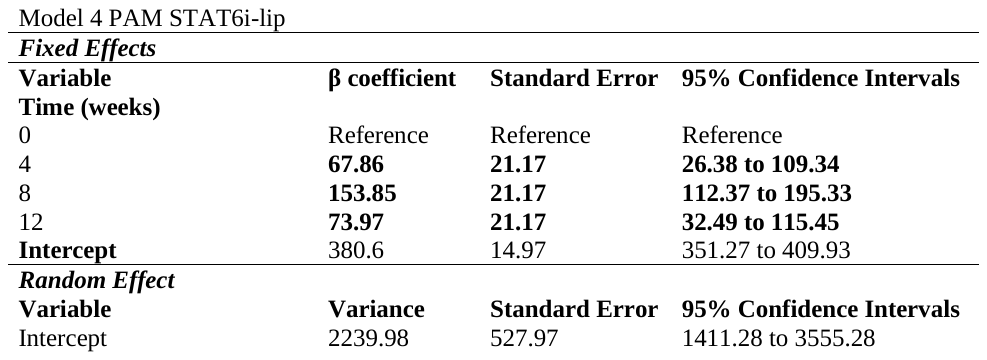

**Supplemental table 6. Individual linear mixed effects models, PAM.** Individual linear mixed effects modelling of pressure pain threshold in Veh-lip **(A)**, Clod-lip **(B)**, STAT1i-lip **(C)**, and STAT6i-lip **(D)**.
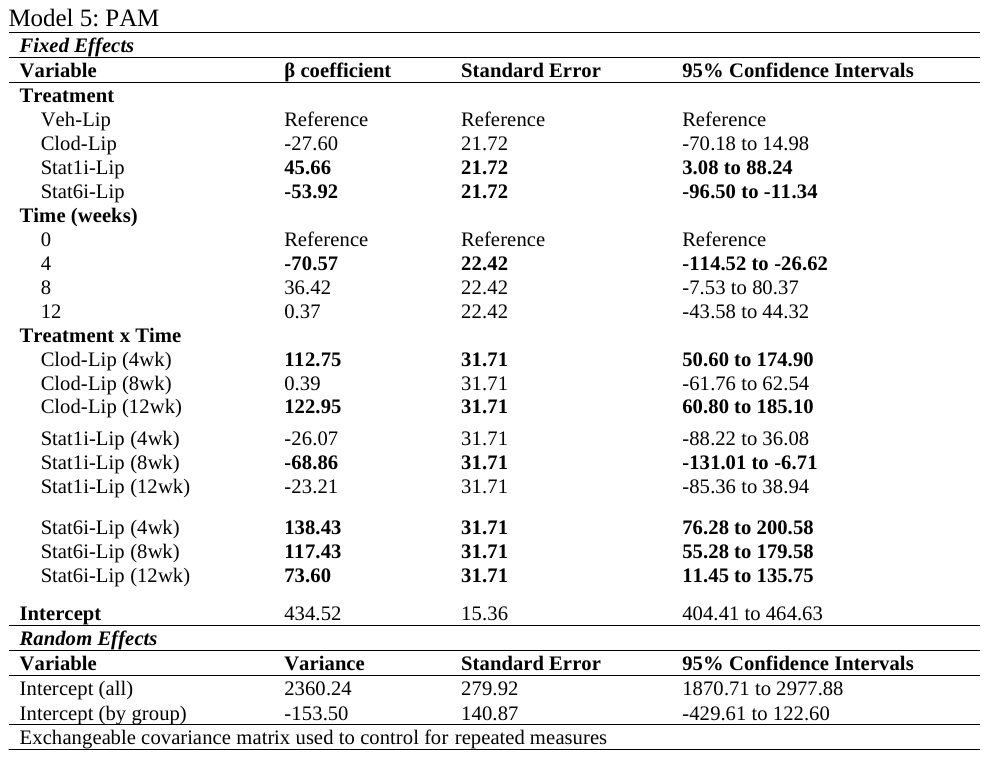

**Supplemental table 7. Linear mixed effects model with pairwise comparisons, PAM.**

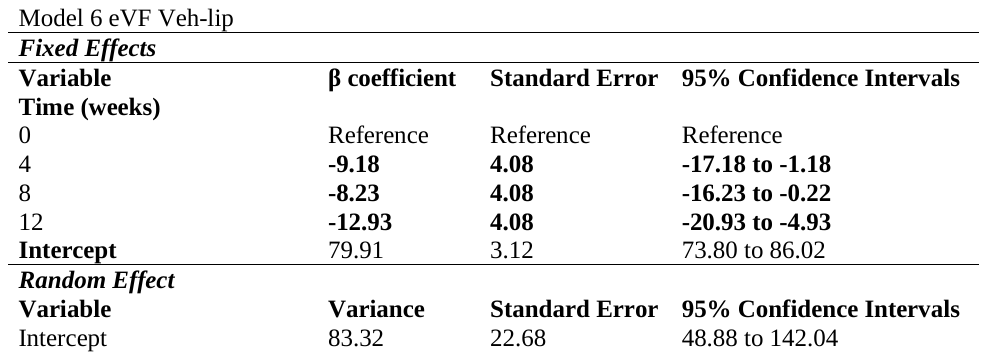

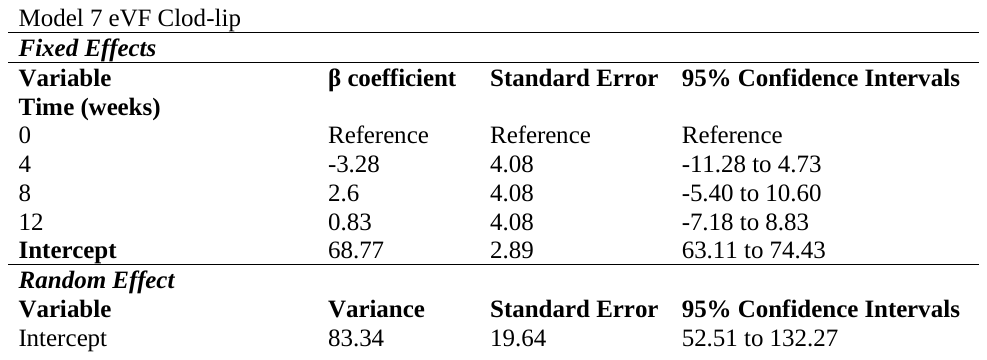

**A**

**B**

**C**

**D**

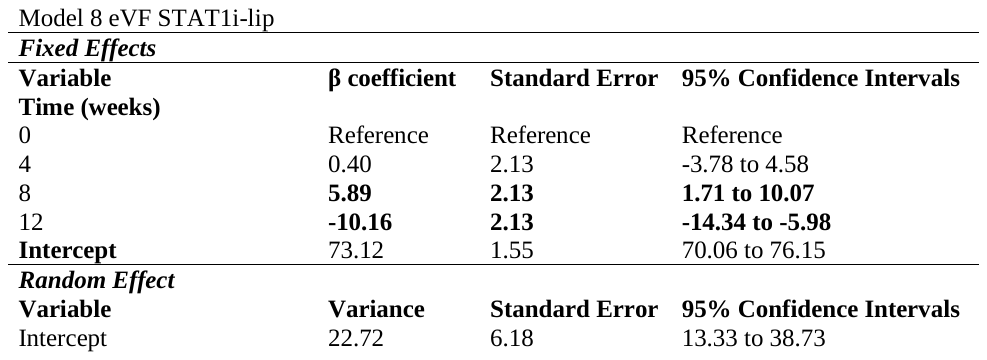

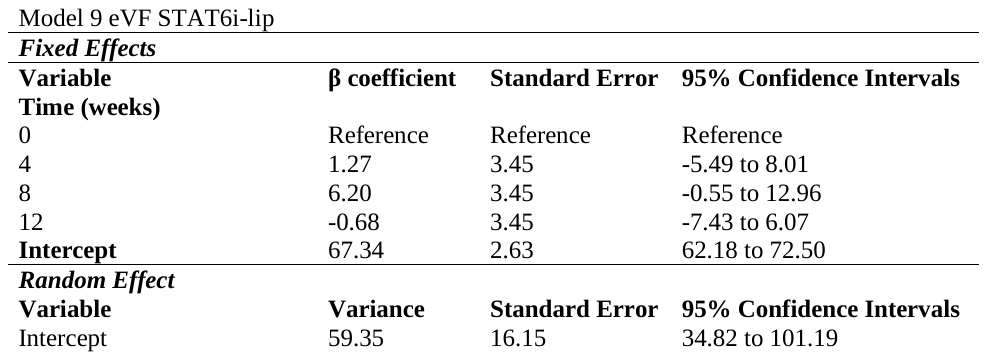

**Supplemental table 8. Individual linear mixed effects models, eVF.** Individual linear mixed effects modelling of hind paw withdrawal threshold in Veh-lip **(A)**, Clod-lip **(B)**, STAT1i-lip **(C)**, and STAT6i-lip **(D)**.

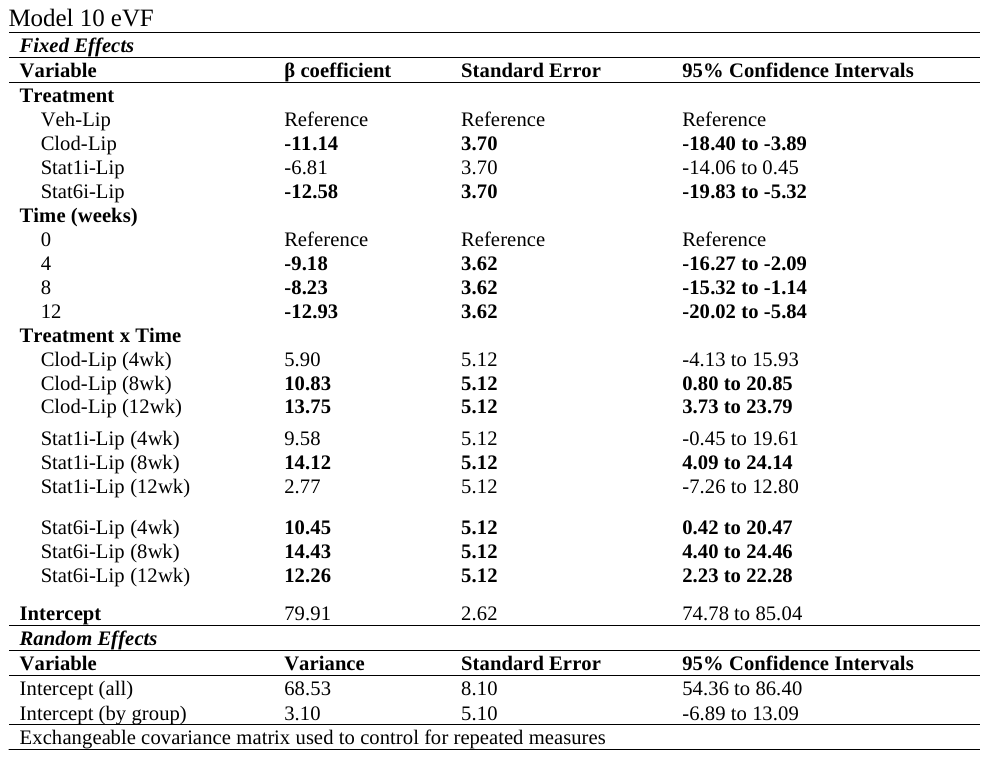

**Supplemental table 9. Linear mixed effects model with pairwise comparisons, eVF.**
